# Rapid Eye Movement Sleep Displays Distinct Fractal Dynamics Between Phasic and Tonic States

**DOI:** 10.1101/2025.04.14.648721

**Authors:** Yiqing Lu, Liang Wu, Jingyu Liu, Yongcheng Li, Yaping Huai

## Abstract

Rapid eye movement (REM) sleep consists of phasic and tonic microstates with unique neurophysiological properties, yet their fractal characteristics remain underexplored. Using Higuchi’s fractal dimension (HFD) analysis of electroencephalographic (EEG) data from healthy adults, this study investigated complexity differences between REM microstates. The results showed that both phasic and tonic REM exhibited significantly lower global HFD values compared to wakefulness, while displaying similar overall complexity levels between microstates. Importantly, phasic REM demonstrated regionally specific reductions in fractal dimensionality, with pronounced decreases observed in frontocentral areas. These localized reductions exhibited a negative association with theta band power, yet remained statistically unrelated to Lempel-Ziv complexity (LZC) measures, indicating that HFD and LZC capture distinct aspects of neural signal organization. The findings reveal that although phasic and tonic REM maintain comparable global complexity, they differ in their spatiotemporal fractal patterns. The association between increased theta power and reduced fractal dimensionality suggests that phasic REM represents a neurophysiological state favoring rhythmic regularity, potentially optimized for internal information processing. These results position HFD as a valuable complementary approach for characterizing REM microstates, with potential applications in elucidating the pathophysiology of sleep disorders.

## 1. Introduction

Sleep is categorized into distinct stages, differentiated by neurophysiological signatures such as characteristic electroencephalographic (EEG) patterns and concomitant autonomic activity (Le Bon 2020). Among these, rapid eye movement (REM) sleep is subdivided into phasic and tonic microstates (phasic REM and tonic REM), each exhibiting unique neural dynamics. Phasic REM is marked by transient events, including ponto-geniculo-occipital (PGO) waves, autonomic fluctuations, and muscle twitches (Dickerson et al. 1993, Fernandez-Mendoza et al. 2009, Gott et al. 2017), whereas tonic REM reflects a stable neurophysiological state with diminished motor activity (Simor et al. 2020). These microstates are not only linked to cognitive functions like memory consolidation (Boyce et al. 2016, Klinzing et al. 2019, van den Berg et al. 2023) and emotional regulation (Wassing et al. 2019, Leitner et al. 2025) but also implicated in disorders such as REM sleep behavior disorder (RBD) and insomnia (Dauvilliers et al. 2018, Feige et al. 2018, Simor et al. 2020, Yang et al. 2024).

Research on REM neurophysiology has employed both linear and nonlinear analytical frameworks. Linear spectral analyses identify distinct EEG power distributions between phasic and tonic REM, demonstrating heightened alpha and beta oscillations during tonic states and increased gamma activity in phasic states (Jouny et al. 2000, Simor et al. 2016). Non-periodic neural activity during REM sleep, distinguished by steeper low-frequency band slopes and flatter high-frequency slopes in phasic states, further provides a novel biomarker for differentiating these states (Rosenblum et al. 2024). Event-related potential (ERP) studies reveal preserved auditory processing during tonic REM in contrast to diminished responses in phasic states (Sallinen et al. 1996, Takahara et al. 2002). Extending these insights, nonlinear approaches such as Lempel-Ziv complexity (LZC) uncover reduced EEG complexity in phasic REM compared to tonic REM, correlating inversely with delta power and positively with alpha activity (Lu et al. 2024). Together, these findings demonstrate distinct information processing mechanisms across REM microstates.

Despite advances in nonlinear EEG characterization, the fractal properties of REM microstates remain underexplored. Fractal analysis, particularly Higuchi’s fractal dimension (HFD), offers a robust framework to quantify signal complexity by assessing self-similarity patterns in time-series data—a feature inherent to nonlinear systems like the brain (Rodriguez-Bermudez and Garcia-Laencina 2015, Lau et al. 2022). Unlike LZC, which evaluates temporal unpredictability through sequence compressibility, HFD captures the multiscale self-similarity and hierarchical structure of neural activity, providing complementary insights into the dynamical organization of sleep stages (Olejarczyk et al. 2022, Armonaite et al. 2023). This approach proves effective in distinguishing pathological from healthy brain states (Gomez et al. 2009, Kawe et al. 2019, Cukic et al. 2020), and may provide novel insights into how phasic and tonic REM contribute to sleep-related disorders.

The differential fractal organization between phasic and tonic REM may reflect distinct modes of consciousness modulation, offering new insights into how hierarchical neural dynamics vary across REM microstates. Such differences in fractal complexity align with evidence from other states of consciousness. For example, fractal dimension measures of brain activity tend to decrease during unconscious states (e.g. NREM sleep or anesthesia) compared to normal wakefulness (Ruiz de Miras et al. 2019), whereas profoundly altered states like those induced by psychedelics show an increase in fractal complexity (Varley et al. 2020). These findings suggest that fractal dynamics sensitively capture the hierarchical neural organization underlying shifts in consciousness, motivating their application to REM sleep microstates. In light of this, the present study employs HFD to quantify the multiscale complexity of phasic and tonic REM sleep EEG signals. HFD was selected over alternative methods due to its superior robustness for short time series and sensitivity to transient complexity fluctuations inherent in brief microstates (Ibáñez-Molina and Iglesias-Parro 2014, Lu and Rodriguez-Larios 2022). Expanding upon our prior LZC analyses, which identified distinct complexity patterns in REM microstates, we now systematically assess fractal dynamics to differentiate phasic and tonic REM, thereby refining their nonlinear neurophysiological profiles. By integrating HFD methodology, this investigation advances the understanding of REM sleep dynamics, offering novel insights into how fractal complexity relates to conscious state modulation during sleep.

## 2. Materials and Methods

### 2.1. Dataset

We analyzed a publicly available REM EEG dataset (https://osf.io/2vptx) originally collected by Simor et al. (Simor et al. 2021). The dataset comprises whole-night polysomnographic recordings from 40 healthy young adults (Study 1: *N*=20, 10 males, 21.7±1.4 years; Study 2: *N*=20, 6 males, 21.6±1.6 years). All participants met strict health criteria and provided written informed consent under local university ethics board approval.

### 2.2. EEG acquisition and preprocessing

For comprehensive methodological details, please refer to reference Simor et al. (2021); a brief summary is provided below. EEG data were acquired using 19 electrodes (10-20 system, Study 1) or 17 electrodes (Fp1/Fp2 were not used, Study 2), mastoids-referenced, with impedances <8 kΩ. Signals were digitized at 4096 Hz and downsampled (Study 1: 1024 Hz; Study 2: 512 Hz). These differences in downsampling rates reflect the original protocols and were preserved to maintain data integrity; however, as all analyses were conducted separately within each dataset using identical procedures, they do not affect the validity of within-study comparisons. Sleep staging followed the American Academy of Sleep Medicine (AASM) criteria, with REM periods classified as phasic (4-second segments with eye movement bursts meeting amplitude criteria) or tonic (4-second segments without notable bursts). In Study 1, each participant contributed 100 segments per state; in Study 2, segment quantities varied for phasic and tonic, but each with a minimum 6-minute duration. For comparisons, we analyzed awake periods (eyes-closed only), including pre-sleep onset wakefulness and post-sleep onset wakefulness when available, applying identical preprocessing pipelines as used for REM segments. Three participants were excluded from Study 2 due to insufficient trials, resulting in final samples of *N*=20 (Study 1) and *N*=17 (Study 2). EEG signals were filtered using a fourth-order Butterworth filter (two-pass, infinite impulse response), with a passband of 0.5-35 Hz. Artifact removal was performed using independent component analyses (ICA) implemented through EEGLAB and FieldTrip toolboxes (Delorme and Makeig 2004, Oostenveld et al. 2011), where 2-4 independent components associated with eye movements were identified based on topography and waveform characteristics. Although Fp1/Fp2 were available only in Study 1, a consistent ICA procedure was applied within each dataset by concatenating phasic, tonic, and wake segments prior to ICA decomposition, allowing for reliable identification of shared artifact components across states.

### 2.3. Estimation of HFD analysis

We quantified EEG signal complexity using HFD, a nonlinear measure that estimates the fractal characteristics of time series data. Following Higuchi’s algorithm (Higuchi 1988), the fractal dimension was derived by calculating the mean curve length of the signal at multiple temporal scales (defined by parameter *k*), where *k* represents the number of intervals used to subsample the time series. The slope of the logarithmic relationship between scale (*k*) and curve length yields the HFD value. For each participant, HFD values were computed across 4-second epochs for each electrode using a maximum scale parameter *k_max_* = 30. This parameter was optimized by identifying the saturation point in HFD values across a range of possible *k_max_* values, ensuring stability in fractal dimension estimation. Condition-specific averages (phasic REM, tonic REM, wakefulness) were obtained by averaging HFD values across epochs within each state. To evaluate the stability of HFD estimates, we performed an additional analysis using a sliding-window approach. HFD was computed within 0.5-second windows, with a step size of 0.25 seconds (i.e., 50% overlap), across each EEG segment for all states. All other parameters remained identical to the main analysis. To assess intra-condition variability in complexity, we calculated the coefficient of variation (CV) of HFD values across all 4-second epochs within each condition, separately for each participant. CV was computed as the ratio of the standard deviation to the mean HFD across segments. The analysis was implemented using custom MATLAB scripts.

### 2.4. Spectral analysis

We examined REM-related EEG spectral changes across all electrodes using Hanning-tapered Fourier transforms with adaptive frequency smoothing (5-cycle windows, 1 Hz resolution). Power was computed for frequency bands: delta (1-4 Hz), theta (4-7 Hz), alpha (8-12 Hz), beta (13-30 Hz), and gamma (31-40 Hz). Spectral analyses were conducted on the same 4-second EEG segments used for HFD estimation, ensuring temporal alignment across analytical procedures. Grand averages were calculated per condition after excluding three participants due to technical artifacts (one in Study 1: anomalous trial concatenation; two in Study 2: extreme delta/theta power outliers), yielding final samples of 19 (Study 1) and 15 (Study 2) participants for spectral and correlation analyses.

### 2.5. Correlation analysis

We computed condition differences (phasic minus tonic REM) for both HFD and spectral power within significant electrode clusters, then assessed their relationship across participants using Pearson’s correlation. Additionally, we examined global HFD-LZC correlations by averaging across all electrodes (phasic minus tonic REM) and performing Pearson’s correlation. The LZC was computed from the same 4-second EEG epochs (Lu et al. 2024). Briefly, each EEG time series was first binarized with respect to its median amplitude, and the LZC index was then obtained by quantifying the number of unique subsequences within the resulting binary sequence, following the standard algorithm proposed by Lempel and Ziv (Lempel and Ziv 1976).

### 2.6. Statistical analysis

HFD values were computed from 4-second EEG epochs using custom MATLAB scripts, averaged per condition (phasic REM, tonic REM, wakefulness) either by electrode or globally. Condition effects were assessed via one-way repeated-measures ANOVA with Greenhouse-Geisser correction when the assumption of sphericity was violated (Mauchly’s test). Post-hoc pairwise comparisons were examined using two-tailed paired *t*-tests, followed by Bonferroni correction.

To address the multiple comparisons problem inherent in EEG data analysis, we implemented a cluster-based non-parametric randomization test (Maris and Oostenveld 2007). The analysis employed a paired-sample design with an alpha level of *α* = 0.05 defining statistical significance. Clusters were formed by aggregating adjacent electrodes demonstrating effects below the predetermined threshold, with cluster-level statistics derived from the summation of *t*-values within each contiguous region. For null hypothesis testing, we performed 5,000 Monte Carlo simulations by randomly shuffling condition labels (phasic vs. tonic REM) to construct an empirical reference distribution of maximum cluster statistics. Final significance determination involved comparing the observed cluster statistics against the 95th percentile of this permuted null distribution, thereby controlling family-wise error rate while maintaining sensitivity to genuine neural effects. Visualizations combined FieldTrip functions with custom scripts.

To complement the cluster-based permutation approach and validate the spatial robustness of the HFD effects, we conducted additional electrode-wise comparisons between states. For each pairwise contrast (phasic vs. tonic REM), we applied the Wilcoxon signed-rank test (two-tailed) at each of the electrodes. The resulting *p*-values were then corrected for multiple comparisons using false discovery rate (FDR) correction (Benjamini and Hochberg 1995), with a significance threshold of 0.05. In addition, all correlation analyses employed FDR correction for multiple comparisons.

## 3. Results

### 3.1. Reduced fractal dimensionality in REM sleep states

A one-way repeated measures ANOVA was conducted to examine differences in HFD across three states: phasic REM sleep, tonic REM sleep, and wakefulness. The HFD values were calculated as the grand average across all EEG channels, representing the overall complexity of neural activity at the whole-brain level. Mauchly’s test indicated violations of the sphericity assumption in both studies (Study 1: *W* = 0.091, *p* < 0.001; Study 2: *W* = 0.224, *p* < 0.001), prompting Greenhouse-Geisser corrections. Both studies revealed strong main effects of state on HFD (Study 1: *F*(1.048, 19.909) = 12.853, *p* = 0.002, *η²* = 0.404; Study 2: *F*(1.126, 21.394) = 46.083, *p* < 0.001, *η²* = 0.708), indicating that neural complexity, as measured by HFD, varied significantly across the different states. Post-hoc analyses employing Bonferroni-corrected pairwise comparisons revealed a highly consistent pattern of results across both studies (Figure 1): no statistically significant differences were observed between phasic and tonic REM sleep states (Study 1: *p* = 0.202; Study 2: *p* = 0.067). However, both phasic and tonic REM sleep showed significantly reduced HFD compared to wakefulness (Study 1: mean differences = -0.041 for phasic REM, -0.037 for tonic REM, both *p* < 0.01; Study 2: mean difference = -0.073 for phasic REM, -0.065 for tonic REM, both *p* < 0.001).

**Figure 1.**
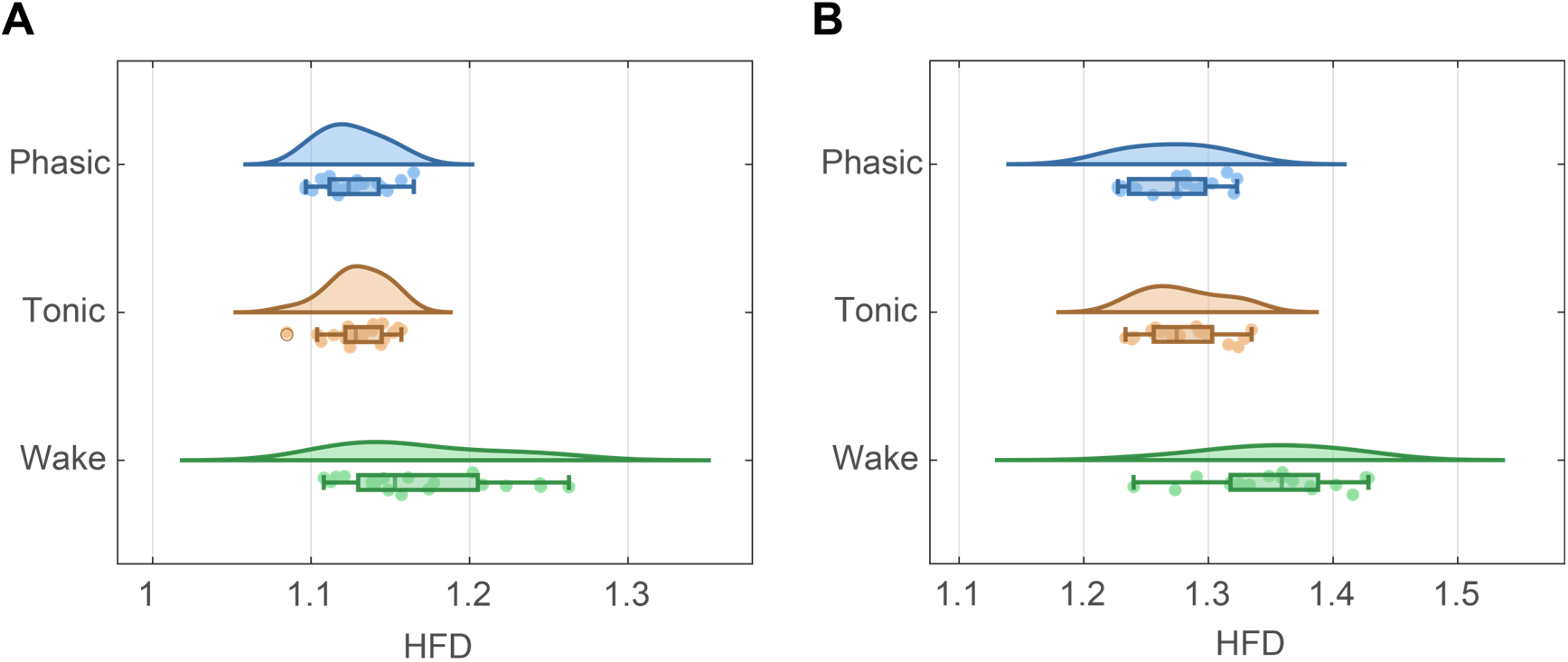
Distribution of individual HFD (averaged across EEG electrodes) across phasic REM, tonic REM, and wakefulness conditions. (A) Study 1; (B) Study 2. For each condition, the density trace shows the smoothed data distribution. The boxplots display the median, interquartile range (IQR), and whiskers (1.5×IQR), with outliers plotted as individual circles when present; individual data points are jittered vertically. Statistical significance was assessed using one-way repeated measures ANOVA with post-hoc test. Both phasic REM sleep and tonic REM sleep demonstrated significantly lower HFD values compared to wakefulness.

### 3.2. Spatial distribution of HFD differences between phasic and tonic REM

Following our initial analysis of grand average HFD across all EEG channels, we next examined the topographic distribution of HFD differences between phasic and tonic REM periods. Cluster-based non-parametric randomization test revealed significantly lower HFD during phasic REM compared to tonic REM (negative clusters, *p* < 0.05; see Figure 2). In Study 1, this reduction was most pronounced (*t*(19) = -20.921, *p* = 0.017) in a frontocentral cluster that included prefrontal, frontal, and central regions (Fp1/Fp2, F3/F4, Fz, C3, Cz). The mean phasic HFD in this cluster (1.128 ± 0.025) was 0.62% lower than during tonic periods (1.135 ± 0.023). Study 2 demonstrated a similar but more extensive pattern of reduced phasic HFD (*t*(16) = -30.471, *p* = 0.008), with a significant cluster distributed across frontal, central, and parietal regions (F7/F8, F3/F4, Fz, C3/C4, Cz, P3/P4). Within this cluster, phasic HFD value (1.263 ± 0.037) was 0.86% lower than tonic value (1.274 ± 0.035), further supporting the robustness of this difference. To assess generalizability, we replicated the topographic analysis using the ANPHY-Sleep dataset (Wei et al. 2024); results similarly showed significantly lower HFD during phasic REM (see Supplementary Figure S1 for details). To confirm the robustness of the observed HFD differences, we conducted additional electrode-wise Wilcoxon signed-rank tests with FDR correction, which largely replicated the frontal-central effects identified by the cluster-based analysis in both studies (see Table S1 for details). Additionally, both phasic and tonic REM exhibited significantly lower HFD than wakefulness across widespread regions (for studies 1 & 2, see Figure S2).

**Figure 2.**
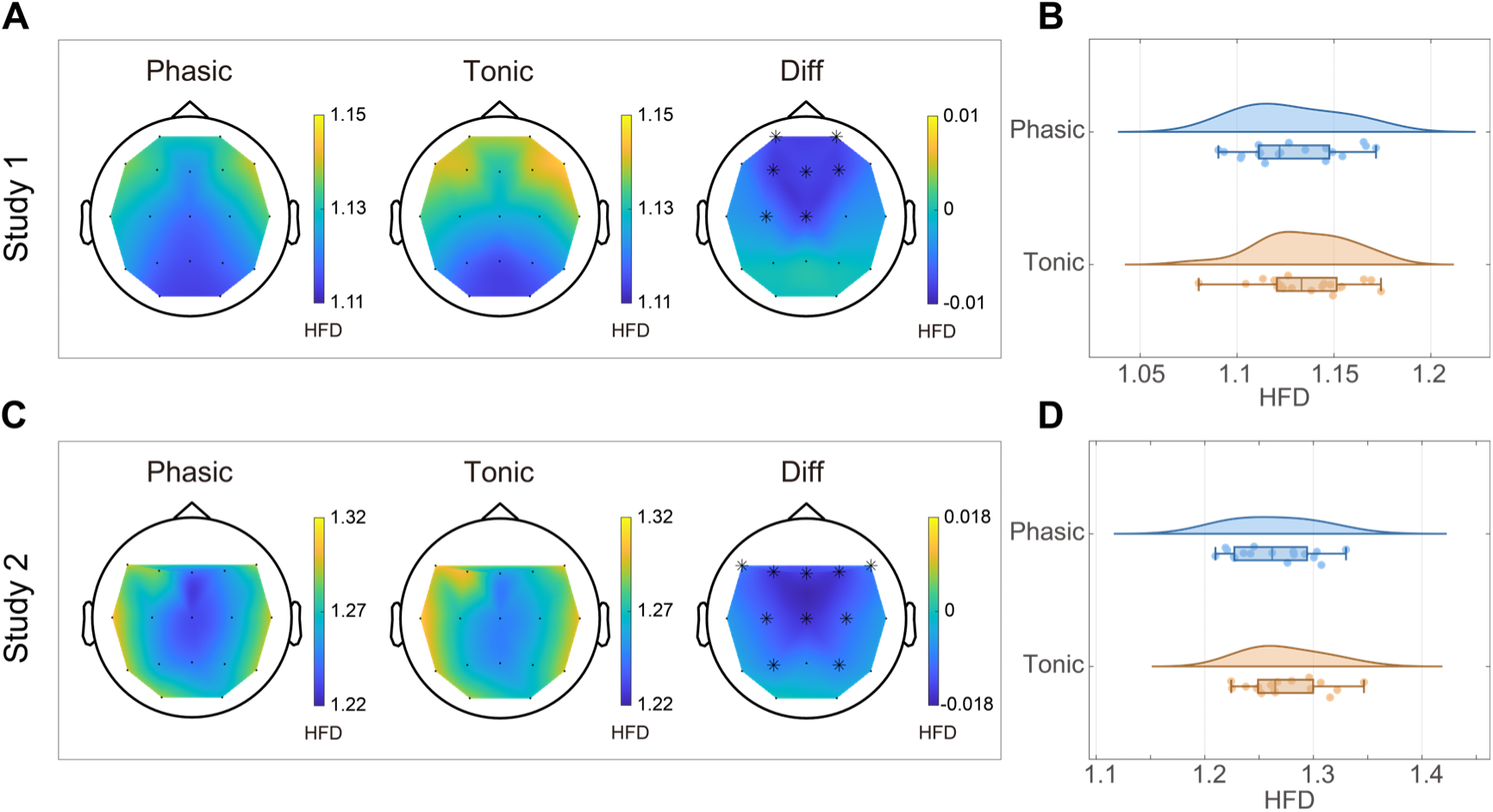
Reduced HFD during phasic REM compared to tonic REM. (A, B) Study 1; (C, D) Study 2. A & C: Topographical maps display mean HFD values per electrode (Fp1 and Fp2 were not recorded in study 2), with difference maps (Diff) highlighting phasic-tonic REM contrasts. Black asterisks denote electrodes with significantly lower HFD in phasic REM (p < 0.05, cluster-based non-parametric randomization test). B & D: Plots of HFD (averaged across significant electrodes) for individual participants (dots), with boxplots indicating median, interquartile range (IQR), and whiskers (1.5×IQR). Density traces illustrate data distributions. In both studies, phasic REM consistently exhibited reduced HFD relative to tonic REM.

A supplementary sliding-window analysis (0.5 s windows with 0.25 s steps) confirmed that phasic REM exhibited significantly lower HFD than tonic REM in both studies, with topographic patterns and numerical distributions closely resembling those observed in the non-overlapping 4-second window analysis (see Figure S3).

### 3.3. Variability of HFD across states

To further characterize the temporal stability of HFD across states, we compared the within-subject coefficient of variation (CV) of HFD values across 4-second segments. Both datasets showed significantly lower CVs in phasic REM compared to tonic REM and wakefulness (Figure S4), indicating reduced intra-state variability during phasic REM (see Supplementary Materials for details).

### 3.4. Correlational analysis of HFD with spectral and complexity measures

Following our examination of grand average HFD and its spatial distribution across phasic and tonic REM periods, we further investigated the functional relationships between HFD and spectral power across multiple frequency bands. Particularly, among all tested bands (delta, theta, alpha, beta, and gamma), only theta power showed a significant negative correlation (FDR corrected) with HFD within the identified electrode clusters. This inverse relationship was consistent across both studies (Study 1: *r* = -0.699, *p* = 9.0×10^-4^; Study 2: *r* = -0.771, *p* = 8.0×10^-4^; see Figure 3), suggesting a specific coupling between decreased signal complexity and increased theta-band activity during phasic REM (relative to tonic REM).

**Figure 3.**
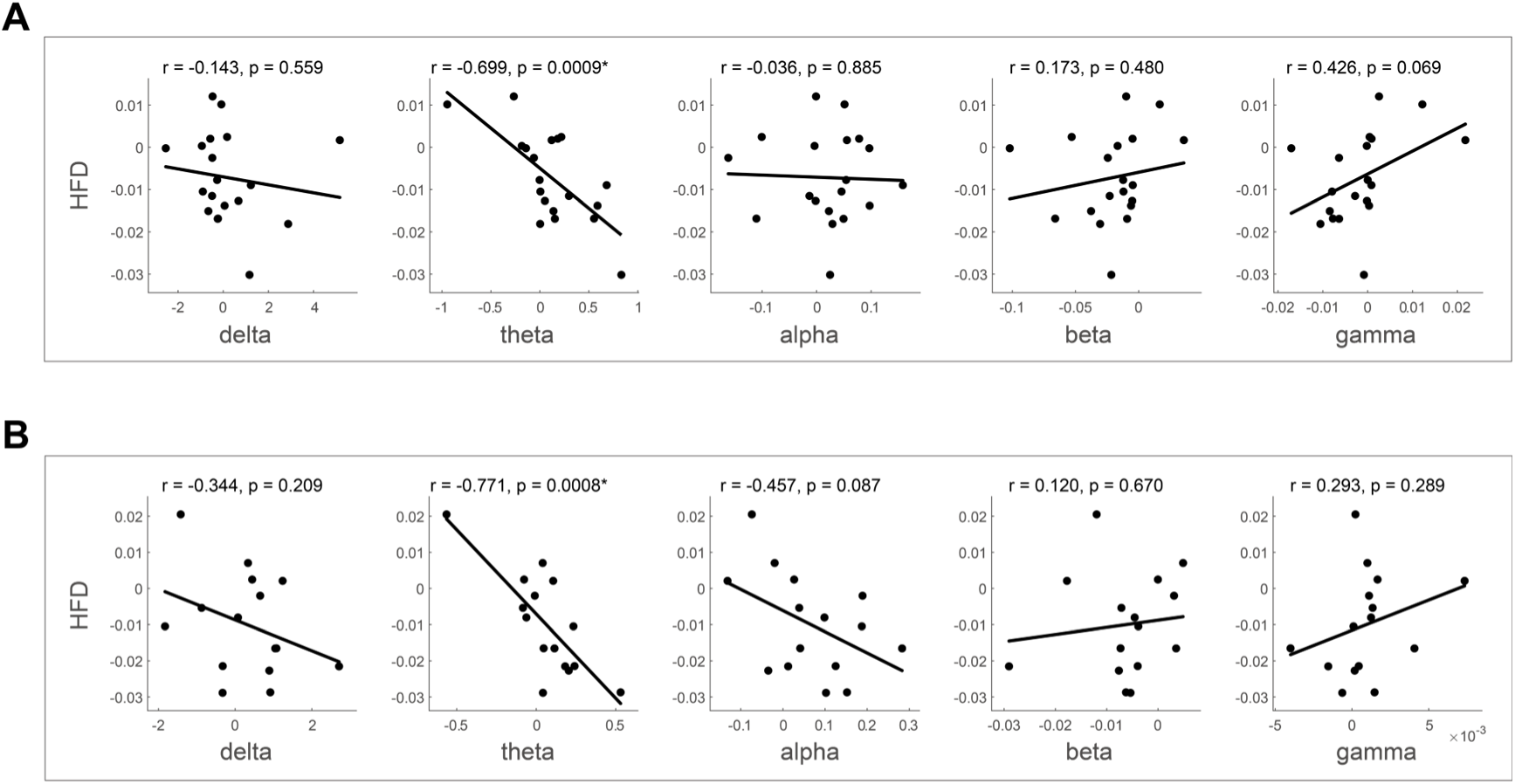
Neurophysiological coupling between HFD and spectral power modulations. (A) Study 1; (B) Study 2. Scatter plots display condition-dependent changes (phasic minus tonic REM) in HFD versus spectral power amplitude for each participant (dots represent electrode cluster averages). Significant negative correlations between HFD and theta power were observed in both studies (Pearson’s r and p-values shown; *p < 0.01, FDR-corrected), indicating that phasic REM-related HFD decreases co-occurred with theta power increases.

Given our prior work demonstrating the relevance of LZC in REM sleep dynamics (Lu et al. 2024), we additionally examined whether the phasic-tonic HFD differences were related to corresponding LZC differences (averaged across all electrodes). However, no significant association emerged (Figure 4), indicating that these complexity measures may capture distinct aspects of neural activity.

**Figure 4.**
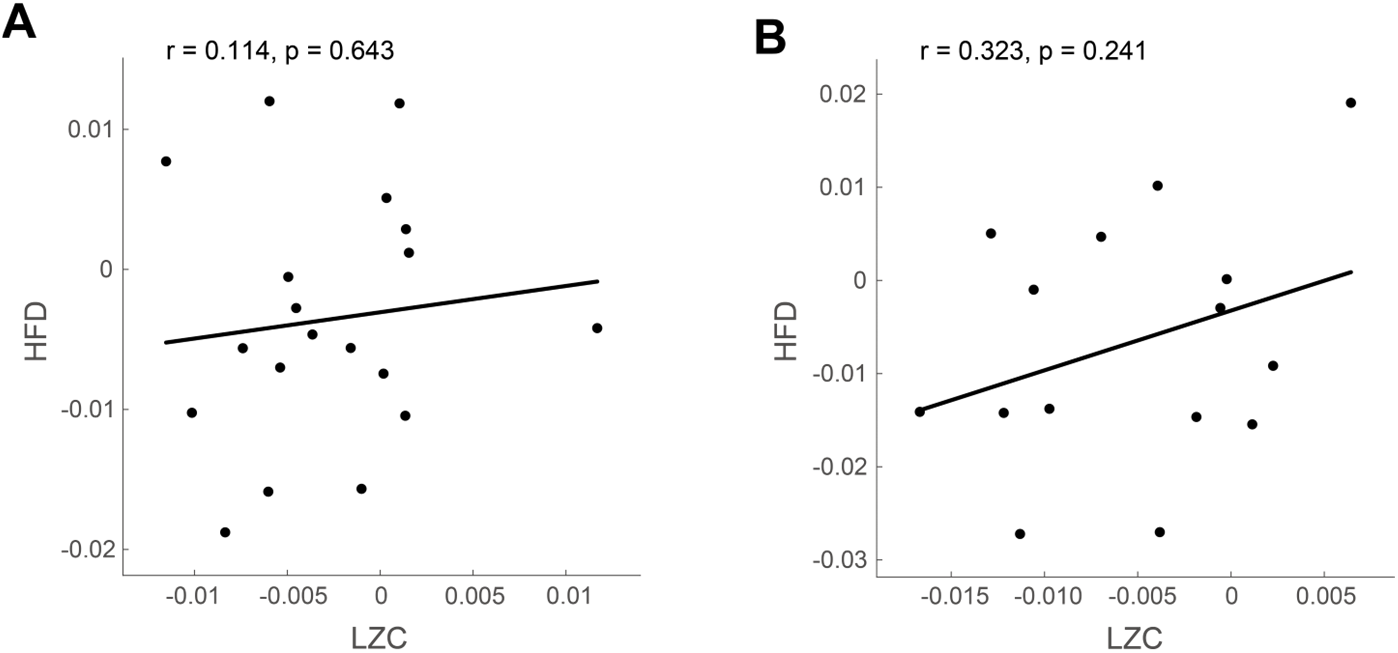
Absence of significant correlation between HFD and LZC. (A) Study 1; (B) Study 2. Scatter plots display condition-dependent changes (phasic minus tonic REM) in HFD versus LZC for each participant (dots represent all electrode averages). No significant correlations were observed between HFD and LZC measures in either study (Pearson’s r and p-values shown; all p>0.05).

## 4. Discussion

The present study provides a comprehensive examination of fractal neural dynamics during REM sleep through HFD analysis, revealing distinct nonlinear EEG characteristics that differentiate phasic and tonic microstates. Our results demonstrate that phasic REM exhibits localized decreases in HFD within frontocentral regions, suggesting microstate-specific modifications in neural signal processing. Both phasic and tonic REM displayed lower fractal dimensions relative to wakefulness, their overall complexity measures did not differ significantly at the global level. Furthermore, we identified an inverse relationship between HFD and theta-band power, independent of LZC measures. These results advance our understanding of REM sleep neurodynamics and highlight the utility of fractal analysis in dissecting neural complexity.

The observed reduction in HFD during REM sleep aligns with prior studies showing diminished nonlinear complexity during sleep compared to wakefulness (Ma et al. 2018, Aamodt et al. 2022, Lu et al. 2024). However, the lack of global HFD differences between phasic and tonic REM diverges from previous LZC results (Lu et al. 2024), suggesting that these metrics assess fundamentally different neural characteristics. This distinction arises from their underlying computational principles: LZC measures temporal unpredictability through sequence compressibility (Lempel and Ziv 1976), while HFD evaluates self-similarity patterns across temporal scales (Higuchi 1988). This divergence implies that REM microstates may share comparable overall complexity but differ in hierarchical signal organization, which is supported by the spatially specific HFD reductions in phasic REM. The frontocentral topography of these differences overlaps with regions implicated in REM-related cognitive processes, including memory consolidation and emotional regulation (van der Helm et al. 2011, Eichenlaub et al. 2018, Kim et al. 2020, van den Berg et al. 2023), suggesting that localized fractal alterations may reflect state-specific neurocomputational demands (Shivabalan et al. 2022, Pakniyat et al. 2024). Moreover, phasic REM also exhibited significantly lower intra-condition variability in HFD, as reflected by reduced within-subject coefficients of variation compared to tonic REM and wakefulness, indicating not only lower complexity but also greater temporal consistency in neural dynamics during this state. The statistical independence of HFD and LZC measurements highlights the complementary nature of these metrics, as LZC primarily reflects temporal unpredictability whereas HFD captures multiscale self-similarity, with their combined use providing a more comprehensive assessment of neural complexity.

The negative correlation between HFD and theta power within phasic REM clusters (compared to tonic REM) provides mechanistic insights into the neurophysiological processes governing REM neurodynamics. Theta oscillations, known to facilitate hippocampal-neocortical communication during memory processing (Boyce et al. 2016), exhibit increased spectral power in phasic REM (Simor et al. 2016). This increased theta activity may drive reduced fractal dimensionality by introducing rhythmic regularity to otherwise irregular neural activity. The rhythmic regularity imposed by theta dominance appears to facilitate the coordination of large-scale neural assemblies for memory reprocessing while promoting perceptual isolation from external inputs through oscillatory entrainment. This sensory disengagement is supported by evidence indicating that phasic REM reflects a neurobiological state preferentially oriented toward internal information processing (Wehrle et al. 2007, Simor et al. 2020), as demonstrated by both attenuated sensory evoked potentials (Sallinen et al. 1996, Takahara et al. 2002) and increased activation of limbic networks (Corsi-Cabrera et al. 2016). The reduction in fractal dimensionality may thus represent an adaptive neural signature of this shifted processing priority, where increased theta synchrony concurrently optimizes memory-related circuit integration and attenuates exteroceptive signaling. These findings extend our understanding of REM sleep by demonstrating how fractal properties and oscillatory dynamics interact to shape both the informational capacity and phenomenological qualities of distinct REM microstates.

Clinically, the differential fractal properties of REM microstates may inform sleep disorder pathophysiology and conscious state modulation. For instance, the frontocentral HFD reductions observed here are consistent with EEG abnormalities reported in RBD (Peng et al. 2021, Valomon et al. 2021), indicating the boundary blurring between dream consciousness and wakefulness characteristic of this disorder. These fractal alterations may represent a characteristic neurophysiological signature of RBD pathology, potentially reflecting impaired gating of conscious experience during REM sleep. The distinct neurophysiological signatures captured by HFD and LZC provide complementary biomarkers that could enhance differential diagnosis. Specifically, HFD’s sensitivity to theta power modulations may prove particularly relevant for disorders involving thalamocortical dysregulation, while LZC’s association with alpha activity, as observed from our previous results, could better characterize disorders of cortical integration (Lu et al. 2024). Future studies should investigate whether these fractal measures can predict conversion from prodromal RBD to neurodegenerative syndromes, quantify consciousness disturbances in REM-related parasomnias, and monitor treatment response in sleep disorders targeting REM mechanisms. Additionally, the development of standardized HFD protocols across clinical populations will be crucial for translating these research findings into practical diagnostic tools.

Several methodological considerations should be acknowledged when interpreting these findings. First, while HFD provides valuable insights into the fractal properties of neural activity, its interpretation remains constrained by the lack of spatial specificity inherent in scalp EEG. Two factors may contribute to slight inter-study variations in HFD topography: (1) electrode coverage differences (Study 2 lacked Fp1/Fp2), which interact with cluster-based statistics by altering adjacent electrode relationships, and (2) moderate sample sizes amplifying individual variability effects. Combining HFD with multimodal neuroimaging techniques could help localize these fractal dynamics to specific neural circuits, thereby clarifying their functional significance. Importantly, our findings maintain methodological robustness through rigorous non-parametric approaches (cluster correction with 5,000 permutations). Second, the restriction to healthy young adults of this study limits the extrapolation of these findings to clinical populations and older individuals, who may exhibit distinct REM sleep characteristics and fractal dynamics. Future studies should include more heterogeneous populations to validate and extend these observations.

In conclusion, this study demonstrates that fractal analysis using HFD provides unique insights into the neurophysiological distinction between phasic and tonic REM sleep. By complementing spectral and complexity metrics, HFD enhances our capacity to characterize the multiscale organization of neural activity during REM sleep, offering new avenues for understanding its role in cognitive processes and sleep-related pathologies.

## Author Contributions

Conceptualization, Y.Lu. and Y.H.; methodology, Y.Lu.; software, Y.Lu. and Y.Li.; validation, Y.Lu., L.W. and J.L.; formal analysis, Y.Lu. and L.W.; investigation, Y.Lu., J.L. and Y.Li.; resources, Y.Lu. and Y.H.; data curation, Y.Lu.; writing—original draft preparation, Y.Lu.; writing—review and editing, Y.Lu. and Y.H.; visualization, Y.Lu.; supervision, Y.Lu. and Y.H.; project administration, Y.Lu. and Y.H.; funding acquisition, Y.Lu., L.W. and Y.H. All authors have read and agreed to the published version of the manuscript.

## Data Availability Statement

The datasets were originally released by Simor et al. (2021, Interoception in REM microstates: osf.io/2vptx) and Wei et al. (2024, ANPHY-Sleep: osf.io/r26fh).

## Acknowledgments

We gratefully acknowledge Simor et al. (2021) and Wei et al. (2024) for making publicly available the EEG datasets used in this research, which were accessed via the Open Science Framework (OSF).

