## Supplementary Materials for "Rapid Eye Movement Sleep Displays Distinct Fractal Dynamics Between Phasic and Tonic States"

### FDR-corrected electrode-wise comparisons

To validate the robustness of the observed HFD differences between states, we conducted additional electrode-wise comparisons (Wilcoxon signed-rank test) using false discovery rate (FDR) correction across all channels. This analysis was performed separately for Study 1 and Study 2. Consistent with the results of the cluster-based non-parametric permutation tests, phasic REM exhibited significantly lower HFD than tonic REM at multiple electrodes after FDR adjustment. The topographically significant electrodes were largely located over frontal and central regions, overlapping with those identified by the cluster-based approach. These findings demonstrate a high degree of convergence between the two statistical methods and support the reliability of the reported HFD effects. The full summary of FDR-adjusted *p*-values is provided in Table S1.

***Table S1.*** *Electrode-wise comparisons of HFD between phasic REM and tonic REM.*

| Electrodes | Study 1 | | Study 2 | |
|---|---|---|---|---|
| | Cluster | Adjusted-*p* (FDR) | Cluster | Adjusted-*p* (FDR) |
| Fp1 | √ | **0.039*** | NaN | NaN |
| Fp2 | √ | **0.039*** | NaN | NaN |
| F7 | - | 0.135 | √ | 0.059 |
| F3 | √ | **0.039*** | √ | **0.024*** |
| Fz | √ | **0.039*** | √ | **0.024*** |
| F4 | √ | **0.039*** | √ | **0.024*** |
| F8 | - | 0.142 | √ | 0.059 |
| T7 | - | 0.353 | - | 0.176 |
| C3 | √ | 0.109 | √ | 0.059 |
| Cz | √ | **0.039*** | √ | **0.024*** |
| C4 | - | 0.340 | √ | 0.059 |
| T8 | - | 0.391 | - | 0.274 |
| P7 | - | 0.970 | - | 0.348 |
| P3 | - | 0.970 | √ | 0.106 |
| Pz | - | 0.970 | - | 0.106 |
| P4 | - | 0.970 | √ | 0.066 |
| P8 | - | 0.970 | - | 0.350 |
| O1 | - | 0.970 | - | 0.906 |
| O2 | - | 0.970 | - | 0.806 |

*Note: Analyses were conducted separately for Study 1 and Study 2. Significant differences (< 0.05) are highlighted in bold. Results confirm the pattern observed in the cluster-based permutation analysis. A checkmark (√) indicates electrodes that belonged to significant clusters (phasic - tonic) in the original cluster-based analysis. Asterisks (*) denote electrodes that remained significant (Wilcoxon signed-rank test) after FDR correction.*

#### Analysis using the ANPHY dataset

To further evaluate the generalizability of our findings, we analyzed an independent open-access dataset (ANPHY-Sleep). For comparability, we selected the same set of EEG electrodes used in the Simor's dataset (Study 1). Among the 29 participants available in the dataset, 7 were excluded due to an insufficient number of phasic REM trials (<30), resulting in a final sample of 22 participants.

A cluster-based non-parametric permutation test revealed significantly lower HFD during phasic REM compared to tonic REM (negative clusters, $p < 0.05$). This effect was most pronounced over a widespread electrode cluster ($t(21) = -81.995$, $p < 0.001$), where mean HFD during phasic REM (1.106 ± 0.019) was 1.16% lower than during tonic REM (1.119 ± 0.019) (see Figure S1). These findings replicate the main effect observed in the original datasets and support the robustness of the phasic–tonic HFD difference across independent datasets.

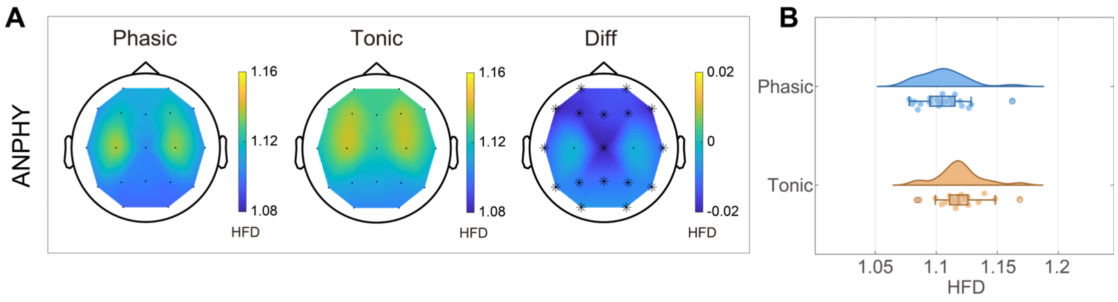


***Figure S1.*** *Reduced HFD during phasic REM compared to tonic REM. (A, B) ANPHY dataset (using 19 electrodes). A: Topographical maps display mean HFD values per electrode, with difference maps (Diff) highlighting phasic-tonic REM contrasts. Black asterisks denote electrodes with significantly lower HFD in phasic REM (p < 0.05, cluster-based non-parametric randomization test). B: Plots of HFD (averaged across significant electrodes) for individual participants (dots), with boxplots indicating median, interquartile range (IQR), and whiskers (1.5×IQR). Density traces illustrate data distributions. Phasic REM exhibited reduced HFD relative to tonic REM.*

### Topographic distribution of HFD differences

Both phasic and tonic REM exhibited significantly lower HFD than wakefulness across widespread regions (see Figure S2).

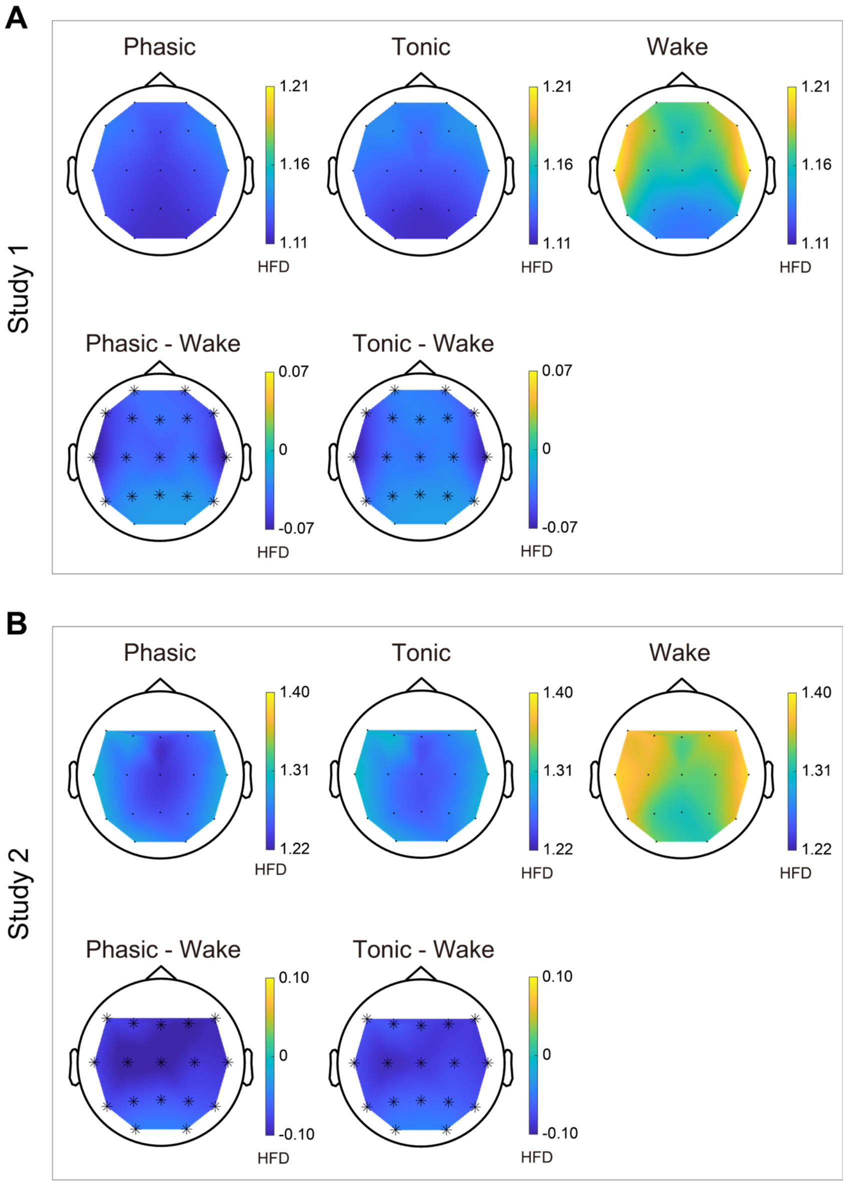


***Figure S2.*** *Reduced HFD during phasic/tonic REM compared to wakefulness. (A) Study 1; (B) Study 2. A & C: Topographical maps display mean HFD values per electrode (Fp1 and Fp2 were not recorded in study 2), with difference maps highlighting phasic-wake REM and tonic-wake REM contrasts. Black asterisks denote electrodes with significantly lower HFD in phasic/tonic REM (p < 0.05, cluster-based non-parametric randomization test).*

### HFD estimates (sliding-window approach)

We also applied a sliding-window analysis followed by cluster-based non-parametric permutation testing. This analysis revealed significantly reduced HFD during phasic REM relative to tonic REM in both datasets (negative clusters, $p < 0.05$; see Figure S3). In Study 1, the effect was localized to a frontocentral cluster encompassing prefrontal, frontal, and central (Fp1/Fp2, F3/F4, Fz, Cz) electrodes ($t(19) = -19.057$, $p = 0.018$), where the mean HFD during phasic REM (1.130 ± 0.025) was 0.70% lower than during tonic REM (1.138 ± 0.023). Study 2 showed a broader cluster extending over frontal, central, and parietal areas (F7/F8, F3/F4, Fz, C3/C4, Cz, P3/P4), with phasic HFD (1.270 ± 0.036) being 0.94% lower than tonic HFD (1.282 ± 0.034), and the cluster-level effect reaching significance ($t(16) = -31.317$, $p = 0.008$). These findings closely replicate the topographic pattern and numerical distributions observed in the original non-overlapping window analysis and further reinforce the reliability of the phasic–tonic HFD difference.

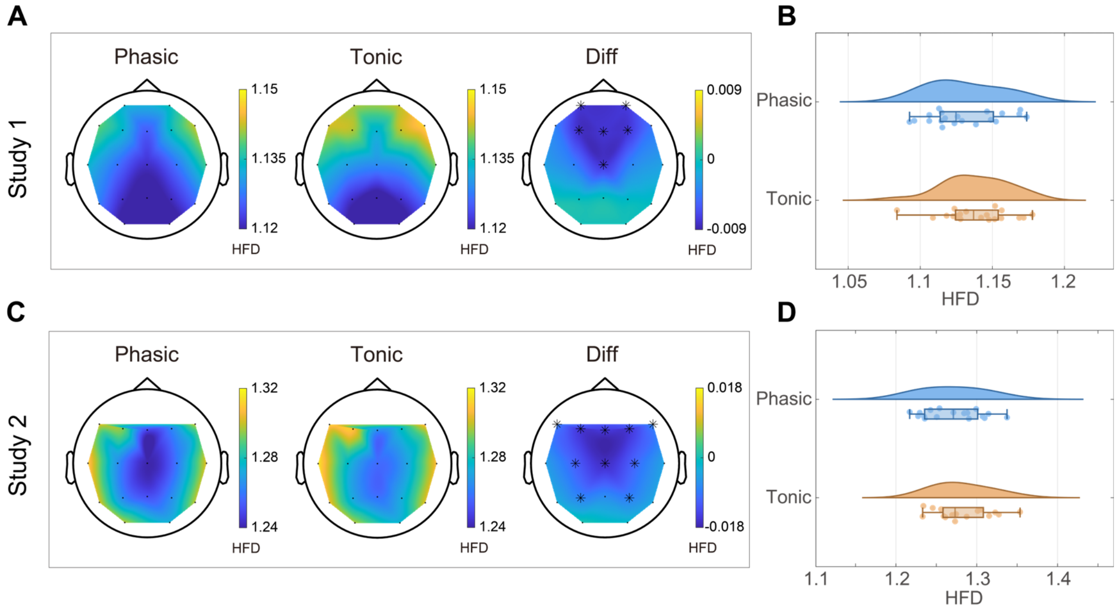


***Figure S3.*** *Reduced HFD (using sliding-window analysis) during phasic REM compared to tonic REM. (A, B) Study 1; (C, D) Study 2. A & C: Topographical maps display mean HFD values per electrode (Fp1 and Fp2 were not recorded in study 2), with difference maps (Diff) highlighting phasic-tonic REM contrasts. Black asterisks denote electrodes with significantly lower HFD in phasic REM (p < 0.05, cluster-based non-parametric randomization test). B & D: Plots of HFD (averaged across significant electrodes) for individual participants (dots), with boxplots indicating median, interquartile range (IQR), and whiskers (1.5×IQR). Density traces illustrate data distributions. In both studies, phasic REM consistently exhibited reduced HFD relative to tonic REM, closely resembling those observed in the non-overlapping 4-second window analysis.*

**Variability of HFD across states**

To assess the variability of HFD across conditions, we calculated the coefficient of variation (CV) of HFD within each state (phasic REM, tonic REM, and wakefulness) for each participant, and conducted one-way repeated measures ANOVAs for Study 1 and Study 2 separately (Figure S4). For Study 1, Mauchly's test confirmed that the assumption of sphericity was met ($W = 0.949$, $p = 0.625$). The ANOVA revealed a significant main effect of state on HFD-CV ($F(2,38) = 9.469$, $p < 0.001$, $\eta^2 = 0.333$). Post-hoc pairwise comparisons (Bonferroni-corrected) showed that phasic REM had significantly lower CVs compared to both tonic REM ($p = 0.006$) and wakefulness ($p = 0.003$), whereas the difference between tonic REM and wakefulness was not significant ($p = 1.000$). For Study 2, Mauchly's test indicated a violation of sphericity ($W = 0.448$, $p = 0.002$), and Greenhouse-Geisser correction was applied. The repeated measures ANOVA again showed a significant main effect of state ($F(1.288,20.615) = 6.442$, $p = 0.014$, $\eta^2 = 0.287$). Bonferroni-corrected comparisons revealed significantly lower CVs in phasic REM compared to tonic REM ($p = 0.005$) and wakefulness ($p = 0.016$), with no significant difference between tonic REM and wakefulness ($p = 0.610$). Together, these findings suggest that phasic REM not only exhibits reduced mean HFD but also demonstrates greater temporal stability in neural complexity, as indexed by lower intra-condition variability.

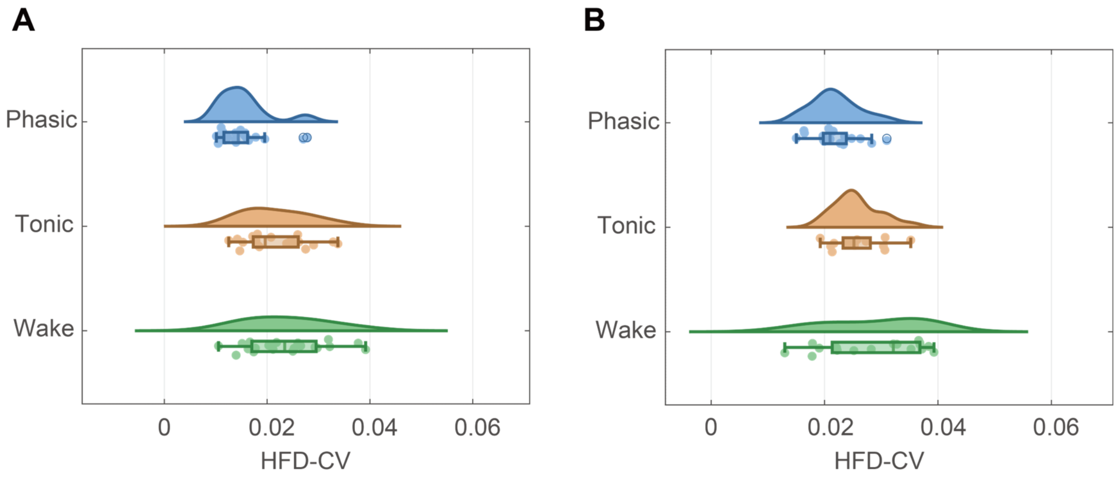


***Figure S4.*** *Distribution of individual HFD-CV (averaged across EEG electrodes) across phasic REM, tonic REM, and wakefulness conditions. (A) Study 1; (B) Study 2. For each condition, the density trace shows the smoothed data distribution. The boxplots display the median, interquartile range (IQR), and whiskers (1.5 ×IQR), with outliers plotted as individual circles when present; individual data points are jittered vertically. Statistical significance was assessed using one-way repeated measures ANOVA with post-hoc test.*
